# The parallel origins of vertebrate venoms

**DOI:** 10.1101/2021.04.26.441528

**Authors:** Agneesh Barua, Ivan Koludarov, Alexander S. Mikheyev

## Abstract

Evolution can occur with surprising predictability when organisms face similar ecological challenges. How and why traits arise repeatedly remains a central question in evolutionary biology, but the complexity of most traits makes it challenging to answer. Reptiles and mammals independently evolved oral venoms consisting of proteinaceous cocktails that allow mapping between genotype and phenotype. Although biochemically similar toxins can occur as major venom components across many taxa, whether these toxins evolved via convergent or parallel processes remains unknown. Most notable among them are kallikrein-like serine proteases that form the core of most vertebrate oral venoms. We used a combination of comparative genomics and phylogenetics to investigate whether serine protease recruitment into the venom occurred independently or in parallel in mammalian and reptilian lineages. Using syntenic relationships between genes flanking known toxins, we traced the origin of kallikreins to a single locus containing one or more nearby paralogous kallikrein-like clusters. Independently, phylogenetic analysis of vertebrate serine proteases revealed that kallikrein-like toxins in mammals and reptiles are homologous. Given the shared regulatory and genetic machinery, these findings suggest a unified model underlying vertebrate venom evolution. Namely, the common tetrapod ancestor possessed salivary genes that were biochemically suitable for envenomation. We term such genes ‘toxipotent’ – in the case of salivary kallikreins they already had potent vasodilatory activity that was weaponized by venomous lineages. Furthermore, the ubiquitous distribution of kallikreins across vertebrates suggests that the evolution of envenomation may be more common than previously recognized, blurring the line between venomous and non-venomous animals.

## Introduction

The extent to which shared history determines repeated evolution of traits remains an important and open question in evolutionary biology. Experiments replaying the tape of life showed that phenotypes can arise through a combination of deterministic forces like natural selection, and stochastic, non-deterministic forces like mutation and genetic drift (Blount, Lenski, and Losos 2018). Selection of homologous and deeply conserved genetic mechanisms can repeatedly produce diverse phenotypes. For example, *hox* genes regulate animal development and are involved in controlling differentiation among body axes, generating the extensive diversity in animal forms (Holland 2013). In plants, modifications of a shared developmental network has repeatedly led to the evolution of bilateral floral symmetry from a radially symmetric ancestor (Citerne et al. 2010). However, most traits are not controlled by master regulators, but emerge from complex interactions within polygenic networks. Yet, how regulatory complexity yields phenotypic novelty remains poorly understood.

To fully realise the course of evolutionary changes, it is essential to have a strong link between genotype and the phenotype they produce (Hoekstra and Coyne 2007; Stern and Orgogozo 2008; Conte et al. 2012). But due to the complex nature of most biological traits, this link is rarely clear. Thus, while short-term evolution via quantitative genetic models is relatively easy to predict, how qualitatively novel traits arise repeatedly is less clear. Exceptionally, reptile and mammalian oral venoms are proteinaceous cocktails where each constituent toxin can be traced to a specific locus, providing an unprecedented level of genetic tractability (Rokyta et al. 2012; Sunagar and Moran 2015; Casewell et al. 2019; Walker 2020). Furthermore, venoms mainly evolve through sequence and gene expression changes of their constituent toxins, providing a strong link between genetic variation and phenotypic change (Sunagar and Moran 2015; Rodríguez de la Vega and Giraud 2016; Safavi-Hemami et al. 2016; Margres et al. 2017; Barua and Mikheyev 2019). Intriguingly, recent work has shown the secretion of diverse and derived toxins by snakes is powered by an ancient conserved gene regulatory network that dates back to amniote ancestors, suggesting a common model for early venom evolution (Barua and Mikheyev 2020). Yet, it is generally believed that individual toxins are convergently recruited, particularly during the evolution of mammalian and reptilian venoms (Schachter 1969; Aminetzach et al. 2009; Casewell et al. 2019). However, this hypothesis has never been explicitly tested.

We tested this hypothesis by examining the evolutionary origins of the most ubiquitous toxins in venom – the serine proteases. Found in all kingdoms of cellular life as well as in viruses, serine proteases are perhaps the most widely distributed group of proteolytic enzymes (Di Cera 2009). Although best characterised in snakes, kallikrein-like (KLK-like) serine proteases are the main components in mammalian venom like that in *Blarina* and *Solenodon*, as well as reptilian venoms in *Heloderma* lizards (Kita et al. 2004; Fry et al. 2010; Casewell et al. 2019). Using a combination of comparative genomics and phylogenetics we were able to resolve the evolutionary origins of venom KLK-like genes. Surprisingly, our results show that mammalian and reptilian serine proteases are homologous and have been recruited into venom in parallel. This implies that the repeated evolution of venom in vertebrates has occurred due to exaptation of already existing components rather than independent evolution of the similar components in different lineages.

## Results

### Genomic organization of the SVL and KLK loci

To determine the genetic history of the venom KLK-like toxins, we identified homologues of the kallikreins in the genomes of mammals, reptiles, amphibians. We specifically focused on tissue kallikreins (TKLs) which are abundant in tissues like pancreas, kidney, as well as in saliva, with functions ranging from mediating blood pressure, muscle contraction, to inflammatory cascades and pain induction (Koumandou and Scorilas 2013). They are also the gene family associated with toxicity of various animal venoms (Fry 2015). Mammalian kallikrein toxins are closely related to the KLK1 gene (Casewell et al. 2019), and we will refer to them as KLK-like toxins. We will refer to their reptilian counterparts as Snake Venom-Like (SVL) toxins.

For example, in humans, TKLs are located in a cluster comprising 15 copies on the 19th chromosome (19q13.4). TKL clusters are also found in other mammalian genomes, though the degree of synteny differs considerably. In venomous mammals like Solenodon and Blarina, the tissue kallikrein loci were modified via tandem duplications of KLK1 and KLK15 genes (Fig 1) (Casewell et al. 2019; Hanf and Chavez 2020). The expanded KLK1 genes contribute to the major toxin component of Solenodon salivary and venomous secretions (Casewell et al. 2019) (Fig 1A). In contrast to the mammals’ single cluster, reptilian genomes have 2-3 gene clusters in the same genomic locus. One of these clusters contains genes that gave rise to viperid snake venom proteases, and has likewise undergone an expansion in this lineage (Fig 1A).

**Figure 1:**
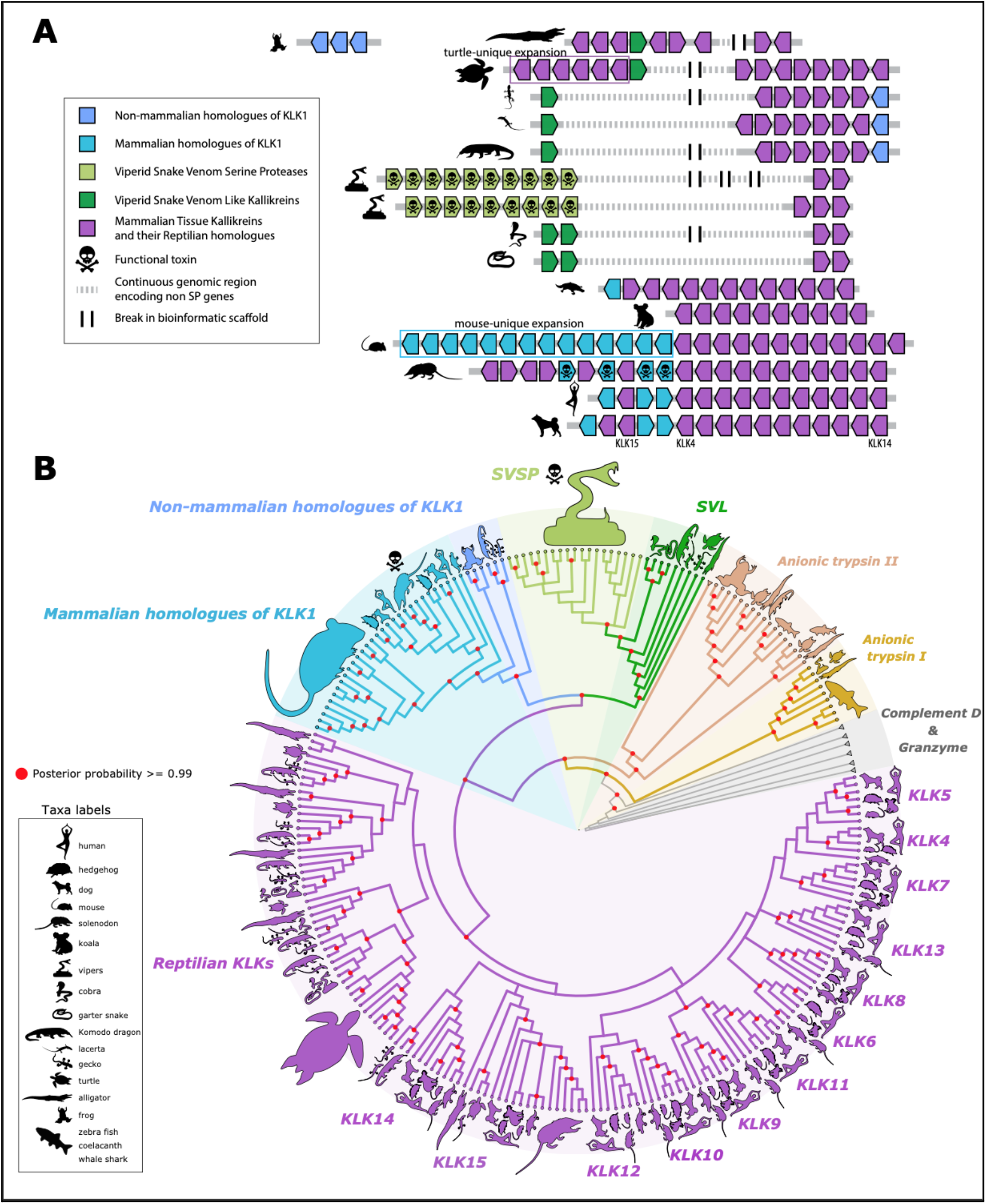
Origins and diversification of tissue kallikreins (TKL) provide insights into the parallel origins of reptilian and mammalian venoms. (A) TKL genes are located at a single genomic locus, and are found in one cluster in mammals and two to three nearby clusters in reptiles. Venom evolution is associated with expansions of toxin-containing gene clusters, but there are also lineage-specific expansions that are not linked to venom evolution (*e.g*., turtles and mice). In existing genomic assemblies the TKL clusters are often fragmented across different scaffolds, but they share many common genes and are clearly contiguous (Supplementary figure 1). (B) Phylogenetic analysis revealed that tetrapod TKLs originated from a common ancestor with vertebrate anionic trypsins, which are commonly expressed in the pancreas and are found elsewhere in the genome. TKLs diverged into two distinct clades, one comprising the KLK4-KLK15 lineages and the other the KLK1/2/3-SVSP/SVL lineage that contains toxipotent genes. Species silhouettes represent members of entire clades rather than a strict node to species demarcation. For a more conventional format please refer to phylogeny (supplementary figure 2 and supplementary dataset 1) in supplementary. Serine protease based toxins are homologs deriving from the same ancestral gene, implying that these toxins originated in parallel venoms in reptiles and mammals.

Reptiles also have other gene clusters that are homologous to mammalian KLK4-KLK14 which are flanked by an ortholog of mammalian KLK15 (with the exception of crocodiles) (Figure 1A). The function of these genes in reptiles is not clear. However, the SVL cluster appears to be a feature unique to reptiles, as it is ubiquitous amongst the examined species of reptile, but is absent from mammalian genomes. In highly venomous snakes like vipers, the expansion of snake venom serine protease (SVSP) genes is linked to the diversification of the venom phenotype (Barua and Mikheyev 2019, 2020), paralleling expansions associated with the evolution of mammalian venoms.

However, we also observed similar expansion of TKL genes in the mouse genome (15 KLK1-like genes) (Karn and Laukaitis 2011) as well as in the Chinese soft-shell turtle (8 copies of TKL genes). The expansion of TKL genes in this manner is highly unusual especially considering both these animals are not venomous. Therefore, this surprising similarity between venomous SVSP, reptilian SVL genes, and venomous KLK1 genes, and non-venomous KLK1 genes suggests some level of shared history.

### Phylogeny of SVL and mammalian KLK genes

We conducted phylogenetic analyses to better understand relationships between and with TKL genes and to identify the likely origin of these genes. Since the TKL-like genes represent a large and diverse gene family it was essential that we sample a wide repertoire of genes across a wide taxonomic distribution. To do this we searched for sequences closely related to KLKs in mammals, reptiles, amphibians, and fish, as classified by NCBI. NCBI’s calculations rely on a combination of calculated orthology and similarity in protein architectures based on sequences in the RefSeq database. This gene set included many non-KLK serine proteases like anionic trypsins, plasminogen, granzyme, and complement B, along with a list of all possible KLK related sequences that are available in NCBI. In order to isolate phylogenetically comparable genes, we used this large gene set (see Methods) as input for OrthoFinder. OrthoFinder classified genes into several large orthogroups. We isolated the orthogroup that contained TKL, SVL, and SVSP genes (supplementary dataset 2) and resolved the phylogenetic relationship between genes within this group. This approach also allowed us to appropriately root our tree and reconstruct the early evolutionary history of TKLs.

Using complement D and granzyme (Fig 1B; grey branches) as outgroups we observed a clear origin of TKLs from two groups of anionic trypsins that are shared between reptiles, amphibians, and fish. After the divergence from anionic trypsins the TKLs split into two separate lineages. While most of the mammalian KLK branching is consistent with previously published mammalian TLK phylogenies (although our tree has better overall support) (Koumandou and Scorilas 2013), we observe several new relationships between genes that were previously not described. First, the SVSP-SVL and mammalian KLK1-KLK2-KLK3 (mKLK1,2,3) genes formed a monophyletic clade sister to the other KLKs (Fig 1b). This topology has high posterior probability (0.99) and was further supported by stepping-stone sampling (Bayes Factor of 111.0 in favor of monophyly between KLK1/2/3 and SVL-like genes, *vs*. the monophy of all KLK-like genes excluding SVSP-like genes). Within the SVL-mKLK1,2,3 clade, the reptilian and mammalian genes form their own sub-clades. The SVL genes appear to group according to the toxicofera classification, with SVL in cobra (*Naja naja*) and garter snake (*Thamnophis sirtalis*) forming a sister clade to the SVSP in elapids and vipers, while non-toxicoferans like the leopard gecko (*Eublepharis macularius*) and the sand lizard (*Lacerta agilis*) forming individual lineages (Fig 1b). Second, KLK15 and KLK14 in reptiles formed a clade with their mammalian homologs, however, several reptile KLKs formed separate reptile specific clades.

### Selection analysis of SVL and mammalian KLK genes

The SVL genes in reptiles are homologous to SVSPs and could have a potential role in imparting toxicity to salivary secretions, as suggested for example in Anguimorph lizards (Fry et al. 2010). Under this assumption, we would expect selection to vary in species believed to have toxic oral secretions, *i.e*. species belonging to the clade Toxicofera, as compared to non-toxicoferans. To test the toxicofera hypothesis we performed branch selection analysis in PAML (Yang 2007). We applied a ‘free ratio’ model for branches leading upto toxicofera and compared its fit to a uniform ‘one ratio’ model for all branches. For a better representation of the toxicofera clade we obtained additional anguimorpha kallikrein sequences from NCBI. We only included coding sequences that encoded for a mature protein and formed a monophyletic clade with our already identified SVL genes (Supplementary figure 3). We did not include venomous snakes in our test because higher selection for toxin genes in venomous snakes is already an established fact and could bias analyses. The two-rate model fits significantly better (LRT, *p* < 0.001) than the uniform one rate model suggesting that toxicoferan SVL genes experienced different selective pressures as compared to non-toxicoferans. We performed the same analysis to test whether venomous mammals experienced different selection as compared to non-venomous ones. The mouse-specific KLK1 genes were not included in this analysis as they are an expansion exclusive to mice and form a clade separate from the other mammalian KLKs including the perceived venomous ones from Solenodon and Blarina. The branches leading up to venomous mammals Solenodon and Blarina experienced selective forces significantly different from the rest of the tree (LRT, *p* < 0.001). While it is difficult to attribute positive selection as the reason for differences in selective pressures from this simple test, some branches (both in toxicofera and venomous mammals) did show high *ω* values (>1) that are indicative of positive diversifying selection (Supplementary dataset 3 and 4). To get a better picture of the selective forces driving evolution of the toxicofera and venomous mammals clade we perform several branch-specific tests using the Datamonkey server (Weaver et al. 2018).

We first used the branch-site unrestricted statistical test for episodic selection (BUSTED) to check for evidence of episodic diversifying election on any site in the gene along any of the branches of toxicofera and venomous mammals (Murrell et al. 2015). For both mammals and reptiles, BUSTED found evidence for diversifying selection in at least one site on at least one test branch (Supp). Since BUSTED revealed joint evidence of branch and site specific selection we used the adaptive branch site random effects model (aBSREL) and mixed effects model of evolution (MEME) to get a better resolution of positive selection in branches of the phylogeny and sites along the gene respectively (Smith et al. 2015; Murrell et al. 2012). Testing the same toxicofera and venomous mammal lineages, aBSREL found evidence for episodic diversifying selection in 1 branch leading to one of the Solenodon KLK1 copies, while in toxicofera it found evidence in 6 branches, one of them leading to the heloderma gilatoxin, another leading to a SVL copy in Haitian giant galliwasp (*Celestus warreni*), and the rest in branches leading upto the radiation of varanids (Fig.2A and B).

**Figure 2:**
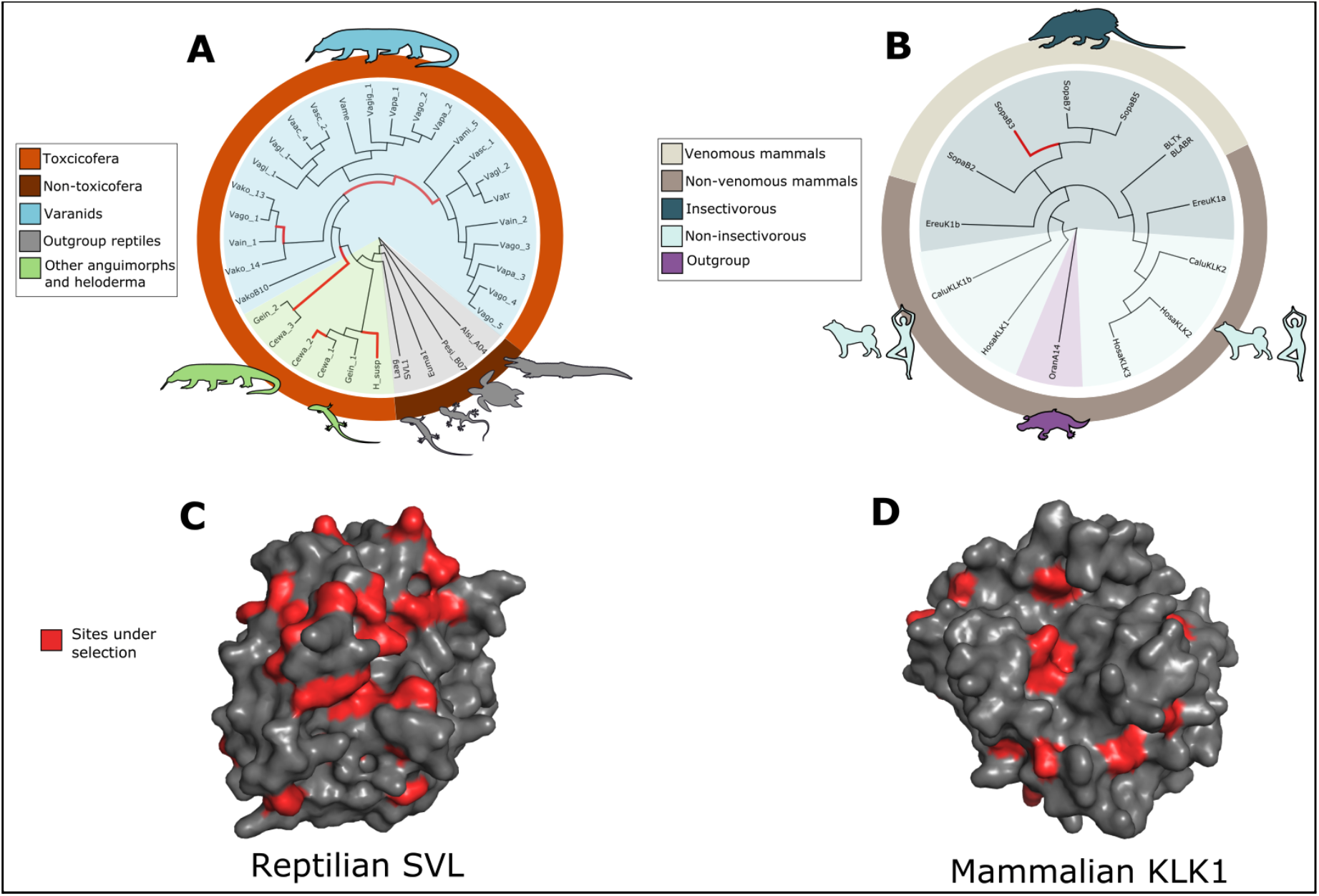
Venomous lineages experienced different selective forces as compared to non-venomous ones for both branch specific and site specific tests for selection.(A) Toxicofera experienced different selection as compared to non-toxicoferan reptiles. aBSREL found evidence for diversifying selection (red branches) in 6 branches within toxicofera. Cewa: *Celestus warreni*, Euma: *Eublepharis macularius*, Gein: *Gerrhonotus infernalis*, H_susp: *Heloderma suspectum*, Laag: *Lacerta agilis*, Pesi: *Pelodiscus sinensis*, Vaac: *Varanus acanthurus*, Vagi: *Varanus gilleni*, Vagl: *Varanus glauerti*, Vagig: *Varanus giganteus*, Vain: *Varanus indicus*, Vako: *Varanus komodoensis*, Vame: *Varanus mertens*, Vami: *Varanus mitchelli*, Vapa: *Varanus panoptes*, Vasc: *Varanus scalaris*. (B) Like in reptiles, venomous mammals experienced different selective pressures as compared to non-venomous mammals. aBSREL found evidence of diversifying selection in only one branch (red) leading to up to a Solenodon copy. Ereu: *Erinaceus europaeus*, Sopa: *Solenodon paradoxus*, BLTx: Blarina toxin, Calu: *Canis lupus*, Oran: *Ornithorhynchus anatinus*. (C) MEME identified 24 sites (in red) in the reptilian SVL that have experienced positive selection. Most of these sites are on the surface. These observations are consistent with previous estimates of high selection on surface residues of toxin serine protease (REF). (D) Unlike reptiles however, only 10 sites on mammalian KLK1s showed evidence of positive selection, with a few on the surface.

The MEME model identified several sites in reptilian SVL genes and mammalian KLK genes that showed significant evidence of positive selection (*p* < 0.05). In reptile SVLs, MEME identified 24 sites experiencing positive selection, while in the mammalian KLKs 10 sites were identified (Fig 2C and D). While some of these sites were in the internal structure of the proteins, the majority of them were on surface residues.

## Discussion

Non-deterministic forces can give rise to evolutionary novelties *de novo*. Several well characterised mechanisms like gene duplication, gene fusion, horizontal gene transfer, etc. are responsible for the birth of new genes (Van Oss and Carvunis 2019). These new genes in turn contribute to species specific processes and generate morphological and physiological diversity (Khalturin et al. 2009). Although non-deterministic processes produce genetic variation (on which natural selection acts), many adaptive traits can be exapted through modifications of already pre-existing characters (Gould and Vrba 1982). Such exaptation has led to the origin of vertebrate oral venoms on at least two levels. Recent work has shown that the ancestral salivary gland gene regulatory mechanisms were exapted in snake venom glands (Barua and Mikheyev 2021). We now show that individual serine protease based toxins used by diverse lineages also evolved from homologous genes. Thus, vertebrate venoms have evolved in parallel, at both the regulatory and also the genetic levels. This suggests that ancient shared history, namely salivary gland regulatory architecture and the presence of homologous genes biochemically suitable for toxicity, have facilitated venom evolution in distantly related taxa.

To determine the role of exaptation in venom evolution, it is important to understand the genetic makeup of adaptive traits, and how they lead to biochemical activity suitable for the envenomation. KLK1 genes in mammals and their reptilian homologs share kininogenase activity, which results in the release of bradykinin, a potent hypotensive agent, when injected into the bloodstream (Komori and Nikai 1998; Kita et al. 2004). This is true even of salivary kallikreins of non-venomous mammals, such as mice, which can induce hypotension and even death (Huang et al. 1977; Hiramatsu, Hatakeyama, and Minami 1980; Dean and Hiramoto 1985). Hypotension is also one of two major strategies which venomous snakes use to immobilize their prey (Aird 2002). The biochemical link between bradykinin-producing enzymes in mammals and snakes was evident to researchers who first characterized kallikrein-like properties of a snake venom enzymes, calling them “the salivary kallikrein of the snake” (Iwanaga et al. 1965). That being said, biochemical similarity does not imply homology. Schachter (1969) wrote in an early review that “kallikreins from different sources are not identical molecules, as originally assumed, nor is it likely that they are derived from a parent molecule”. The belief that kallikreins in mammals and reptiles have different origins continued to the present day (Aminetzach et al. 2009; Casewell et al. 2019). Our analysis overturns the assumption that held for over 50 years, and shows that genetic mechanisms underlying venom evolution are actually homologous. Indeed, all KLK1- and SVL-like genes share a common origin at the dawn of the tetrapods when they formed nearby gene clusters (Fig 1A). However, even from within this family of paralogous proteases, venoms evolved from more closely related homologous genes (Fig 1B).

### Evolution of tetrapod venoms by kallikrein exaptation

Most exaptations have bifunctional intermediates where both the old and new functions are preserved (Manley, Fay, and Popper 2004; Luo 2007). This bifunctional nature likely allows for a gradual transition from one phenotypic state to another. This is the standard model of snake toxin evolution, which presuppose gene duplication prior to the acquisition of novel function (toxicity) (Fry, Vidal, et al. 2009; Hargreaves, Swain, Hegarty, et al. 2014). This is indeed observed in a recent study reconstructing the evolution of metalloproteinase toxins, which evolved from adam28 disintegrin by duplication and modification, such as the loss of a transmembrane domain improving solubility (Giorgianni et al. 2020). However, it appears that kallikreins already possess biochemical activity suitable for envenomation (vasodilation via bradykinin production); we have called such genes ‘toxipotent’. Interestingly, serine protease genes in viperid snake venoms have undergone extensive duplication, with no clear distinction between an ancestral gene and its derived toxic counterparts, which is at odds with the classical venom evolutionary model. So far genomes of venomous mammals and heloderma lizards are insufficiently well characterized to see whether this has been the case as well.

Barua and Mikheyev (2021) proposed a unified model of early venom evolution in mammals and reptiles, suggesting that venoms evolved when kallikreins already present in saliva increased (via higher copy number) and became more effective (via sequence level changes). In this study we were able to reconstruct the evolution of ubiquitous kallikrein-based toxins via phylogenetics based on extensive taxonomic sampling, and gene orthologs accurately selected from the wide range of serine proteases found in the genome based on phylogenetic and syntenic proximity. First, we found that copy number changes accompany the evolution of venom (*e.g*., snakes and Solenodon), but some lineages experience copy number expansions without evolving venom (mice and turtles, Fig. 1A). Second, we found that venomous taxa (gila monster and Solenodon) indeed have a higher rate of nonsynonymous changes in the rates of venom evolution, consistent with selection for novel function (Figure 2). Intriguingly, we also find evidence of selection in reptilian members of the Toxicofera clade, such as varanid lizards, where the existence of venom is debated (Figure 2A) [(Fry, Wroe, et al. 2009; Koludarov et al. 2017) though see (Hargreaves, Swain, Logan, et al. 2014; Sweet 2016; Borek and Charlton 2015)]. However, the presence of toxipotent genes in the saliva of many animals makes the line between venomous and non-venomous animals less clear. Indeed, there could be many taxa that lie on the continuum between what we currently perceive as venomous and non-venomous, as most tetrapods already possess the requisite machinery for venom evolution.

## Conclusion

In this study we expanded our knowledge on the phylogeny of kallikreins (KLKs), and for the first time resolved the origin of tissue kallikreins (TKLKs) in tetrapods. The tetrapod lineage of TKLKs evolved from an ancient serine protease that gave rise to vertebrate anionic trypsins. From here the tetrapod TKLKs diverged into the KLK4-KLK15 group and the toxicopotent KLK1-SVL-SVSP lineage. These toxicopotent homologs eventually diversified and became a part of venom in snakes, some lizards, as well as some shrews and Solenodon. We overturn a decades-long assumption that venoms originate through a combination of constraint and convergence, and show that shared history and parallelism can explain the repeated evolution of toxins in venoms. Parallelism is sometimes considered a process that led to the rise of phenotypic similarity in closely related species (Conte et al. 2012). While this perspective can account for a shared molecular basis and history, the numerous exceptions to this prevents it from being definitive (Steiner et al. 2008; Manceau et al. 2010). It is more appropriate to consider parallelism as the use of shared molecular mechanisms to produce convergent phenotypes, irrespective of their taxonomic proximity (Rosenblum, Parent, and Brandt 2014). We illustrate this by showing that venom in mammals and reptiles originated multiple times in parallel by modifying the same gene family despite 300 million years separating these lineages. Thus ancient conserved molecular mechanisms can continue to be a source of adaptive novelty, allowing nature to replay the tape of life, albeit with a new perspective.

## Methods

### Genomic analysis

We used publicly available vertebrate genomes of good quality (Supplementary table 2) to establish location and synteny of the Kallikrein clusters. We used genomes for which RNA-seq verified genomic annotations were available as a reference point and created an extensive map of the genes that flank SVL and TKL in those genomes (Supplementary table 2). That allowed us to establish syntenic relationships of those regions in different genomes. We then proceeded to use those flanking genes as a database to BLAST (NCBI-BLAST v.2.7.1+ suite, blastn, e-value cutoff of 0.05, default restrictions on word count and gaps) the rest of the genomes that were less well annotated. That gave us a number of genomic scaffolds that potentially contained KLK genes. We used those for the second round of BLAST (tblastx, e-value cutoff of 0.01) against a database of exons extracted from well-annotated mammalian TKL and viper SVL genes. Positive hits were checked by eye in Geneious v11 (https://www.geneious.com) and any complete exons were manually annotated and later merged into CDS of newly annotated genes if the exon order and count was in accordance with existing reliable KLK annotations. All resulting genes that produced viable mature peptides were then used for the phylogenetic analysis.

### Phylogenetic analysis

All viable genes located in the previous step were translated into proteins and aligned with selected publicly available sequences of interest using L-INS-i method of MAFFT software v7.305 (Katoh and Standley 2013) with 1000 iterations (--localpair --maxiterate 1000). These parameters were used for all subsequent alignments. The publicly available serine protease sequences were obtained from NCBI. Using human KLK1 (gene ID: 3816) as a search query we obtained a list of all similar genes that were estimated based on synteny information and conserved protein domains. Full list of genes in NCBI can be found [here](https://www.ncbi.nlm.nih.gov/gene/3816/ortholog/similargenes/?scope=33208). We selected sequences from Human (*Homo sapiens*), mouse (*Mus musculus*), dog (*Canis lupus familiaris*), hedgehog (*Erinaceus europaeus*), Lacerta (*Lacerta agilis*), garter snake (*Thamnophis elegans*), habu (*Protobothrops mucrosquamatus*), Chinese soft-shell turtle (*Pelodiscus sinensis*), alligator (*Alligator sinensis*), frog (*Xenopus tropicalis*), zebra fish (*Danio rerio*) coelacanth (*Latimeria chalumnae*), and whale shark (*Rhincodon typus*). Alignments were refined by hand using Geneious v11 (https://www.geneious.com) to make sure that obviously homologous parts of the molecule (like the cysteine backbone) were aligned properly. A final alignment with 50% masked gaps was used to make the tree (Supplementary dataset 2). We carried out phylogenetic analysis using MrBayes (v3.2.3) (Ronquist et al. 2012). The analysis used a mixed amino acid model and was carried out across two parallel runs for 200 million generations (Altekar et al. 2004), by which point the standard deviation of split frequencies reached 0.0065. Half of the trees were removed as burn-in and the rest summarized to compute posterior probabilities. We also computed Bayes factor support for monophy of SVSPs and KLK1/2/3 *vs*. the monophyly of all KLK genes by stepping-stone sampling of tree space with corresponding backbone constraints for 50 million generations (Xie et al. 2011).

### Selection analysis

Alignments for sequence analysis were carried out using MAFFT alignment tool, implementing the E-INS-i algorithm with BLOSUM62 as the scoring matrix (Katoh and Standley 2013). All alignments were trimmed to remove signal peptide. The phylogeny was constructed based on the relationship of genes obtained from the phylogenetic analysis of genes (Fig 1B). Additional anguimorpha kallikrein can be found in (Supplementary dataset 3). To test for selection on branches leading to venomous animals we used maximum likelihood models implemented in CodeML of the PAML package (Yang 2007). The log likelihood was compared between test branches (venomous animals) vs background branches (non-venomous animals), and significant difference in models was determined using a log likelihood ratio test (Supplementary dataset 4). Tests for adaptive evolution using BUSTED, aBSREL, and MEME analysis were carried out on the Datamonkey server (Weaver et al. 2018). The three-dimensional protein models for SVL and KLK1 were generated using a homology search implemented on the Phyre2 server (Kelley et al. 2015). PyMOL was used for visualization (PyMOL Molecular Graphics System, Schrödinger, LLC).

## Author contributions

I.K. and A.S.M conceptualised the study. I.K. annotated sequences and performed synteny analysis. All three authors analyzed the data and wrote the manuscript.

